# Myosin-dependent partitioning of junctional Prickle2 toward the anterior vertex during planar polarization of Xenopus neuroectoderm

**DOI:** 10.1101/2022.08.26.505384

**Authors:** Chih-Wen Chu, Carsten Stuckenholz, Mona Gerges, Lance A. Davidson

## Abstract

Planar cell polarity (PCP) of tissues is established by mutually exclusive partitioning of transmembrane proteins Frizzled and Vangl with their respective binding partners, Dishevelled and Prickle. While the amplification and maintenance of this pattern have been well studied, it remains unclear how the anterior-biased protein localization is initiated. Moreover, PCP protein complexes are located at adherens junctions and their polarization requires the activity of non-muscle myosin II (NMII), but how NMII contributes to PCP is not fully understood. Here we analyze time-lapse images of mNeonGreen-tagged Prickle2 (Pk2) in mid-gastrula stage *Xenopus* presumptive neuroectoderm and demonstrate that Pk2 puncta move along bicellular apical junctions in a biased manner toward the anterior vertex, where the Vangl-Pk complexes are normally enriched. In addition, length changes of bicellular junction segments flanking each Pk2 punctum are often different from each other, and appear more dynamic near the vertices, suggesting that Pk2 movement is driven by intrinsic junction heterogeneity. Reducing NMII activity eliminates the anterior movement, and surprisingly, increases the displacement of Pk2 puncta. By assessing the correlation between Pk2 movement and the relative positioning of each Pk2 punctum along apical junctions, we uncovered that NMII activity is required for the anterior Pk2 movement by maintaining the elongation of posterior junction segment while inhibiting Pk2 movement toward both vertices flanking the junctions. Importantly, we found that the tight junction protein ZO-1 partitions alongside Pk2. Our findings provide the first evidence of biased partitioning of junctional PCP proteins toward the anterior vertex and support the hypothesis that NMII activity facilitates Pk2 polarization not via a direct transport but by regulating intrinsic dynamics of the bicellular junction.

## Introduction

Epithelial tissues in multicellular organisms often establish planar cell polarity or PCP, i. e. the global alignment of the polarity of individual cells, in the tissue plane during development. PCP and its downstream pathways control many morphogenetic events such as convergent extension, apical constriction, and ciliogenesis (Adler, 1992; Butler and Wallingford, 2017; Peng and Axelrod, 2012; Roszko et al., 2009). At the molecular level, PCP is established by mutual exclusion of two “core” protein complexes toward opposite poles within each cell. Asymmetric distribution of the transmembrane protein Frizzled (Fzd), Van Gogh-like (Vangl), and Flamingo/Celsr within a cell is amplified by clustering the same kind of protein complex while antagonizing clusters of the other, and this process is mediated by their respective cytoplasmic binding partners Dishevelled (Dvl; Dsh in *Drosophila*) and Prickle (Pk) (Butler and Wallingford, 2017; Peng and Axelrod, 2012; Strutt and Strutt, 2009).

The initial asymmetry of PCP components is generated by polarity cues in the environment, but exactly how this asymmetry is created is not fully understood. Several studies have demonstrated selective stabilization of PCP proteins at apical junctions in a context-dependent manner (Aw et al., 2016; Butler and Wallingford, 2018; Chien et al., 2015; Newman-Smith et al., 2015; Ossipova et al., 2015). Of note, live imaging has shown that junction shrinkage and rearrangement can create asymmetric distribution of PCP proteins (Aw et al., 2016; Butler and Wallingford, 2018), providing direct evidence of PCP establishment via junction remodeling. In addition, PCP components can be delivered to cell junctions via microtubules and F-actin cables aligned with the axis of polarity cues (Blair et al., 2006; Ikawa and Sugimura, 2018; Luxenburg et al., 2015; Mahaffey et al., 2013; Matis et al., 2014; Ren et al., 2007; Shimada et al., 2006). Furthermore, PCP components themselves are well known to regulate the cytoskeleton via RhoGTPases (Schlessinger et al., 2009), as PCP signaling alters levels of actomyosin contractility and enables emergent polarity of cortical actomyosin movements (Kim and Davidson, 2011). However, in many cases the symmetry breaking of PCP components has not been well documented, potentially due to difficulties to perform live imaging while planar polarization is taking place and the likely feedback mechanisms that strengthen planar polarity.

Core PCP proteins are known to cluster at apical junctions (Butler and Wallingford, 2018; Cho et al., 2015; Ossipova et al., 2015; Strutt et al., 2016; Strutt et al., 2011). In *Xenopus* neuroectodermal cells undergoing convergent extension, Vangl-Pk clusters are often asymmetrically distributed along individual adherens junctions and enriched near the anterior vertex (Butler and Wallingford, 2018; Ossipova et al., 2015); yet, the underlying mechanism responsible for their asymmetry remains elusive. Molecular mechanisms for PCP localization are likely to involve cortical actomyosin and cell-cell adhesions that are ubiquitous within apical junctional complexes. Recent studies have revealed that individual junctions, i.e. segments between vertices, exhibit mechanical heterogeneity accompanied by asymmetrically distributed cadherin cis-clusters (Huebner et al., 2021; Strale et al., 2015). Moreover, it is known that contractility of junction-associated F-actin by non-muscle myosin II (NMII) can drive junction segment length change (Hara, 2017; Vicente-Manzanares et al., 2009), and that contraction of sub-domains within junction segments by RhoA can translate ZO-1 clusters along junctions (Stephenson et al., 2019). These observations suggest that junction heterogeneity, i.e. the uneven distribution of tension within an individual junction segment, is responsible for asymmetric localization of proteins along junctions. However, while there are many studies regarding protein movement within a 2-D plane of plasma membrane (Jacobson et al., 2019), the movement of proteins along apical junctions has not been reported.

In *Xenopus* embryos, PCP localization has been reported in vivo by the late gastrula or early neurula stage (Chien et al., 2015; Ossipova et al., 2015), suggesting that PCP is established during gastrulation. To better understand this process, we performed live imaging using mNeonGreen-tagged Prickle 2 protein (Pk2) to track the location of the Vangl-Pk complex at apical junctions of presumptive neuroectodermal cells. The high fluorescence efficiency of mNeonGreen allowed low expression that minimally impacted PCP polarity and yet still allowed highly resolved imaging of Pk2 within the apical junctional complex. Our analyses of time-lapse sequences revealed that Pk2 accumulates in discrete puncta at the junction and that these puncta translate along junctions with a bias toward anterior cell vertices. Consistent with the requirement of NMII activity for PCP in the neural plate (Ossipova et al., 2015), reduction of NMII activity inhibited the anterior bias of Pk2 puncta movement but surprisingly increased Pk2 puncta velocity. In addition, Pk2 puncta within the same junction do not move uniformly, reflecting intrinsic junction heterogeneity. Together, our data suggest that anterior accumulation of Pk2 reflects heterogeneous dynamics within bicellular junctions and that NMII contributes to Pk2 polarization by regulating Pk2 puncta movements within the apical junction complex.

## Results

### Pk2 puncta translate along apical junctions toward the cellular vertex

To better understand how the apical distribution of Pk2 becomes polarized, we carried out live imaging of presumptive neuroectoderm expressing mNeonGreen (mNG)-Pk2 during gastrulation. Consistent with the distribution of GFP-Pk2 (Butler and Wallingford, 2018), mNG-Pk2 became enriched at anterior cell boundaries in neuroectoderm at as early as late gastrula (Figure 1- Figure Supplement 1, stage 11.5, top panels). Surprisingly, we found that Pk2 signals at the apical junction often appeared as puncta and that these puncta translated directionally along bicellular junction segments toward the anterior vertex (Figure 1A and 1B, Figure 1- Supplemental Movie 1), raising the possibility that this movement contributes to Pk2 polarization.

**Figure 1.**
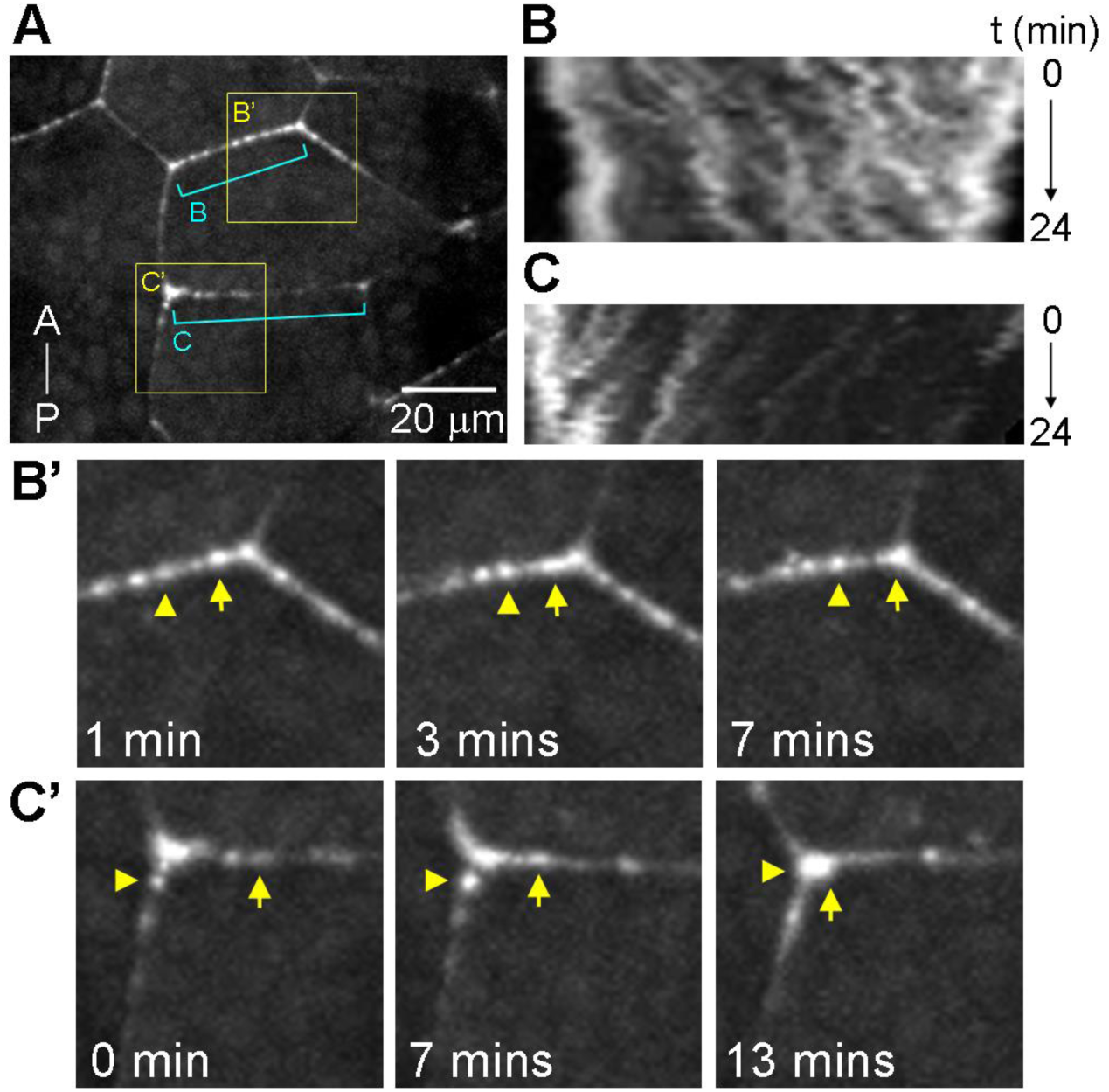
Pk2 movement along the apical junctions. Presumptive neuroectoderm of embryos expressing mNG-Pk2 was excised at stage 11 and imaged. (A) A still of a representative movie at t = 0 min. The Antero (A)-posterior (P) axis is shown. (B) (C) Kymographs of junction B and C in (A), respectively. (B’) (C’) Stills of the magnified view of box B’ and C’ in (A), respectively. Note the movement of Pk2 signals (arrowheads and arrows) toward the cell vertex over time.

To test this possibility, we sought an unbiased method to quantify Pk2 puncta movements along junctions. Apical junctions are acutely responsive to local and global cues, leading to cell shape changes and network remodeling. Normal patterning and mechanical processes as well as perturbations may produce heterogeneous dynamics within apical junction segments and vertices in single cells. To reduce potential bias, e.g. a priori selection of junction segments or cases where tissue mechanics or morphogenesis is globally disrupted, we first established a pipeline for image acquisition and analysis of Pk2 within apical junctions (Figure 2A). Embryos injected with mRNAs encoding mNG-Pk2 and membrane-mCherry (mem-mCherry, a membrane-targeted fluorescent protein) were raised until stage 11, and the presumptive neuroectoderm was dissected as a dorsal isolate (Davidson, 2022) and subjected to live imaging under a spinning disk confocal microscope. Morphogenesis of the dorsal isolate, an organotypic explant of the dorsal anlagen, preserves both local cell behaviors and long range tissue rearrangements but constrains motions to a 2D plane allowing stable imaging of the apical surface of the neurectoderm. Time-lapse image sequences of 10 minutes were acquired and segmented using the intensity of mem-mCherry (Aigouy et al., 2010). Pk2 intensity profiles along segmented junctions between two vertices were computationally straightened to produce kymographs, which were then subject to additional analysis using machine learning algorithms (Jakobs et al., 2019) to extract trajectories of Pk2 puncta. Trajectories were manually corrected when they were visibly mis-connected. Puncta velocity and directionality were calculated from individual trajectories using custom software. Since dorsal tissues undergo convergent extension at this stage (Keller and Danilchik, 1988), individual junction lengths vary during the time course of image acquisition. Changes in junction length can confound analysis of sub-junctional dynamics and limit the reliability of Pk2 puncta absolute velocities as an indicator of whether a punctum actually translates directionally along a junction. To overcome this limitation, we defined the “relative displacement” of each Pk2 punctum as its start-to-end relative position change times the starting junction length (Figure 2B). We additionally calculated a “normalized velocity (NV)” for each punctum by dividing the relative displacement by the duration of their track. Velocities were defined as positive for puncta moving anteriorly and as negative for puncta moving posteriorly based on the position of junction segment vertices at the start of each time-lapse sequence, and the absolute value of velocity of NV reflects the magnitude without directionality information.

**Figure 2.**
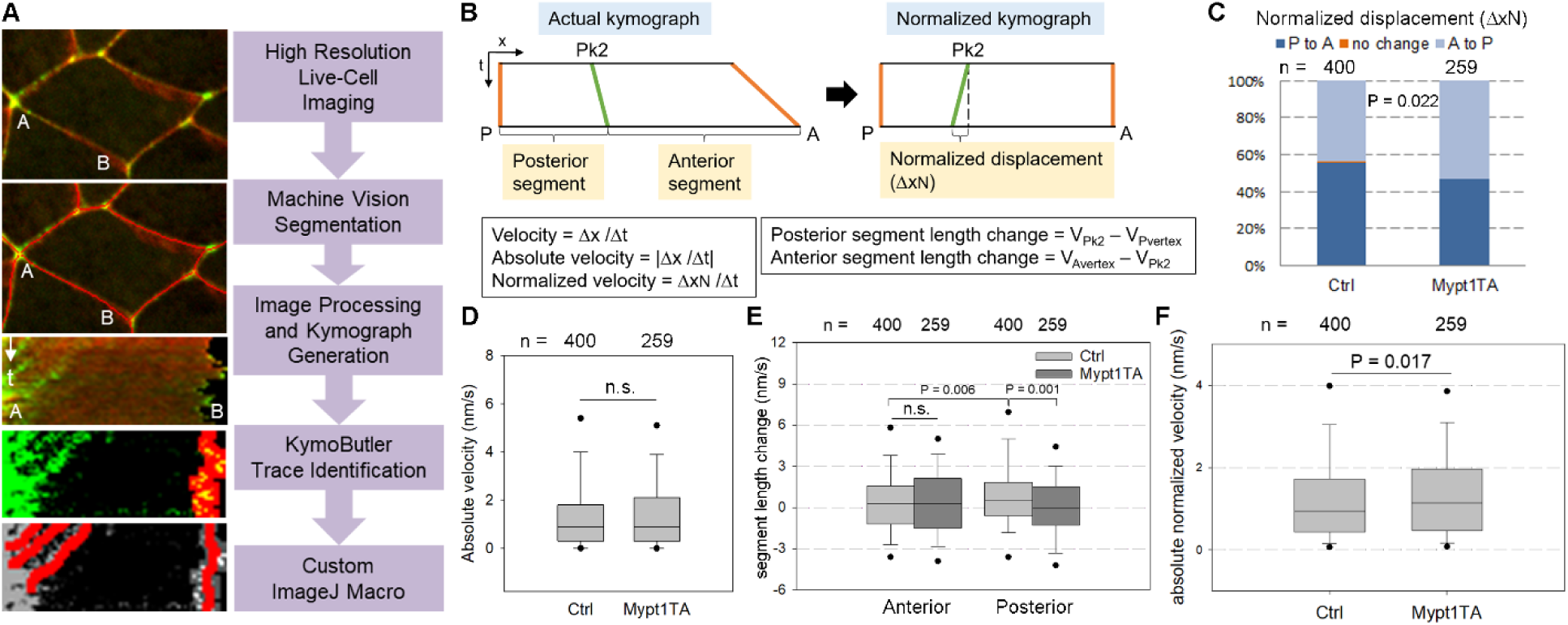
NMII activity is required for the anterior movement of Pk2 along the junctions. (A) Image analysis pipeline. (B) Schematic illustration of Pk2 movement parameters. Kymographs are normalized based on the relative position of Pk2 puncta at each time point. (C) Distribution of normalized Pk2 moving direction in control embryos and embryos expressing Mypt1TA. n, number of puncta. Chi-square test was performed to obtain the *p* value. (D-F) Box plots showing the 25th, median, and 75th percentiles of the data. The error bars represent the 10^th^ and 90^th^ percentiles, and the dots mark the 5^th^ and 95^th^ percentiles. n.s., not significant. (D) Absolute velocity of Pk2 puncta in control and Mypt1TA-expressing embryos. Mann-Whitney rank sum test was performed to obtain the *p* values. (E) Length change of the segments anterior or posterior to Pk2 puncta in control and Mypt1TA-expressing embryos. A positive difference means the corresponding junction segment is lengthened. Two-way ANOVA was performed to obtain the *p* values. (F) Absolute normalized velocity of Pk2 puncta in control and Mypt1TA-expressing embryos. Mann-Whitney rank sum test was performed to obtain the *p* values.

### Pk2 puncta translocate bi-directionally along junction segments, but there is a bias toward the anterior vertex that depends on non-muscle myosin II

To test whether this method can quantify the gradual polarization of Pk2 puncta to the anterior side, we analyzed 400 puncta movements from 68 junctions in 4 embryos from 2 clutches and found that 56% of Pk2 puncta moved toward the anterior vertex, 43% moved toward the posterior vertex, and 1% showed no movement (Figure 2C), demonstrating the efficacy of our method. Furthermore, the anterior-ward bias of Pk2 puncta movement supports previous observations of Pk2 localization at anterior vertices. To understand whether this bias is associated with Pk2 polarization, we next sought to perform the same analysis in tissues with defective planar polarization and see whether Pk2 movement is still biased or not. We disrupted Pk2 polarization in embryos by expressing the constitutively type 1 mutant of Mypt1 (Mypt1TA), the myosin-binding subunit of myosin II phosphatase (Weiser et al., 2009). This mutant protein reduces NMII activity by dephosphorylating myosin light chain, and it has been shown to disrupt Vangl2 polarization in the neural plate (Ossipova et al., 2015). Low doses of Mypt1TA mRNAs were injected unilaterally into embryos, so that PCP was disrupted locally but gastrulation of the whole embryo was largely unaffected. As expected, Mypt1TA expression inhibited mNG-Pk2 polarization in neuroectoderm in late gastrula (Figure 1- Figure Supplement 1A-C; Figure 1-Supplemental Movie 2). Pk2 puncta in junctions shared by two Mypt1TA-expressing cells were tracked, and we found that only 47% of Pk2 puncta moved to the anterior vertex in the presence of Mypt1TA, which is significantly lower than that in the control embryos (Figure 2C). This data suggest that NMII activity can contribute to PCP formation by biasing Pk2 movement along the junction. Curious about the distribution of NMII activity during Pk2 polarization, we assessed the localization of SF9, a live probe for NMII (Hashimoto et al., 2015; Nizak et al., 2003; Vielemeyer et al., 2010), and found that it is located at the cell junctions without apparent polarization (Figure 2- Figure Supplement 1). We decided to look further into the characteristics of junctional Pk2 movement and how it is regulated by NMII activity.

### Non-muscle myosin II regulates junction elongation from the posterior vertex and slows Pk2 translocation

From these results we suspected the NMII could play a direct role in transporting Pk2 puncta. We next calculated the start-to-end velocity of Pk2 puncta movement and found most puncta translated at velocities less than 2 nm/sec (Figure 2D). This velocity is considerably slower than most of the known cytoskeleton motors (Milo et al., 2010) (as an example, myosin-mediated vesicle transport has a speed of 200 nm/sec), and close observation did not reveal transient rapid movements commonly observed during motor-mediated vesicle transport. Such a low velocity argues against motor-mediated transport of puncta, and our analyses of the diffusion coefficient and moment scaling spectrum (MSS) slope of Pk2 puncta suggest that at least 68% of the puncta move via sub-diffusion processes (Sbalzarini and Koumoutsakos, 2005)(Ferrari et al., 2001) (Figure 2- Figure Supplement 2). Furthermore, expression of Mypt1TA had no effect on Pk2 absolute velocity (Figure 2D). This suggests that Pk2 translation is not due to directional transport as a cargo by myosin II but rather reflects (1) puncta diffusion near the plasma membrane or within the cell cortex, or (2) type 2 movement in junctional complexes as junctions undergo heterogeneous growth and shrinkage. To assess the possibility that junction length is regulated differentially on both sides of Pk2 puncta, we tracked the change of distance between each punctum and their anterior or posterior vertex (referred to as the anterior or posterior segment, respectively; Figure 2B) as segment length change. Given that translation of Pk2 puncta is biased towards the anterior end of the bicellular junction, we expected the anterior junction segments to be shortened (negative segment length change), or the posterior segments to be lengthened (positive segment length change), or both if the punctum was undergoing diffusion. Indeed, the posterior segment length change in control junctions was mostly positive, while the anterior segment length change did not have a strong bias (Figure 2E). Interestingly, expression of Mypt1TA specifically diminished the positive change of posterior segment length (Figure 2E), suggesting that NMII activity is required for the biased lengthening of posterior segments.

Time-lapse sequences revealed that the distribution of positive and negative Pk2 NV in control junctions was not significantly different from those in junctions of cells expressing Mypt1TA (Figure 2- Figure Supplement 3); however, Pk2 puncta overall translate faster in Mypt1TA-expressing cells (Figure 2F; absolute magnitude of NV), indicating that NMII activity limits Pk2 translocation. In light of the potential effect of NMII activity on both the directionality and magnitude of junctional Pk2 movement, we decided to focus our analysis on the magnitude of NV and frequency of anterior movements of the Pk2 puncta.

### Non-muscle myosin II limits Pk2 puncta movement toward the nearest vertex

Since posterior segments of bicellular junctions lengthen in a NMII-dependent manner (Figure 2E), we analyzed the potential correlation between the starting position of the puncta along the junction and their subsequent movement. If the growth of posterior junction segment occurs at a specific location in the junction, e.g. at the vertex versus the middle of the segment, Pk2 puncta located posterior to that location will not translate anteriorly. In junctions of control cells, we found no correlation between the punctum’s starting position and either the absolute NV or direction of motion (Figure 3A and 3B). To our surprise, expression of Mypt1TA allowed Pk2 puncta to move toward the nearest vertex, eliminating the anterior bias in puncta motion (Figure 3A). Moreover, Pk2 puncta at the central region of Mypt1TA-expressing cell junctions translate more rapidly, having a significantly higher absolute NV compared to puncta at other regions or puncta at the same region of control junctions (Figure 3B). Taken together, we find that junctional Pk2 puncta move anteriorly regardless of their relative position and that NMII prevents Pk2 puncta from moving indiscriminately toward the nearest vertex.

**Figure 3.**
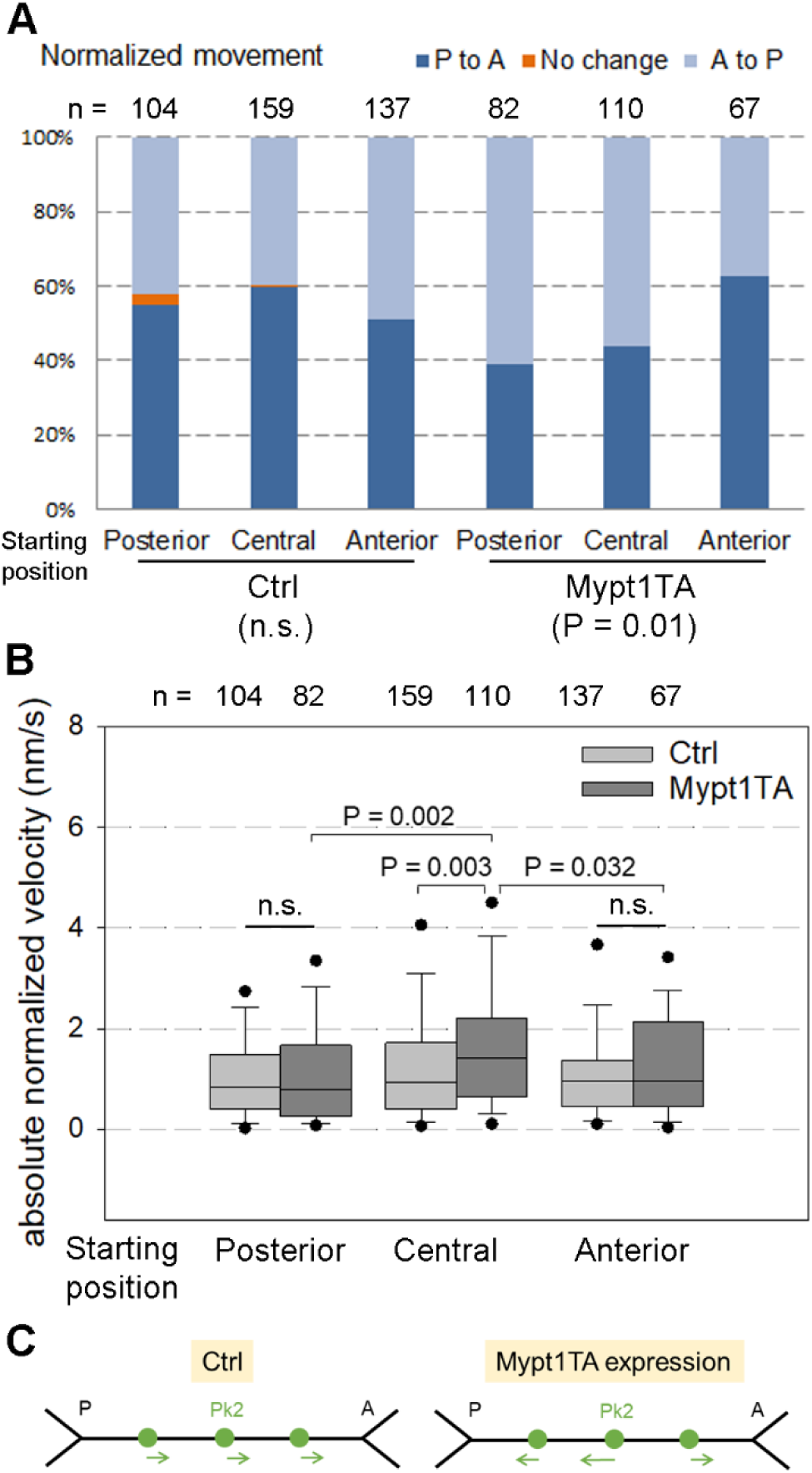
NMII activity is required to restrict Pk2 movement. (A) Distribution of normalized Pk2 moving direction based on the starting position of Pk2 puncta at junctions in control and Mypt1TA-expressing embryos. Chi-square test was performed to obtain the *p* values. (B) Absolute normalized velocity of Pk2 puncta based on their starting position at junctions in control and Mypt1TA-expressing embryos. Two-way ANOVA was performed to obtain the *p* values. (C) Summary of the finding. Expression of Mypt1TA disrupts anterior-ward Pk2 movement by allowing Pk2 movement toward both vertices and increasing Pk2 velocity.

Our finding that Pk2 puncta move toward the nearest vertex when NMII contractility was reduced indicate that Pk2 puncta within a junction can move in opposite directions. Interestingly, about 62% of junctions in control cells exhibited mixed directionality of Pk2 puncta movement, whereas only 40% of junctions in Mypt1TA-expressing cells did (Figure 4A). While this result could be biased by the spatial distribution of Pk2 puncta in each junction, it suggests that Pk2 puncta within a single junction often do not move in the same direction due to junction heterogeneity.

**Figure 4.**
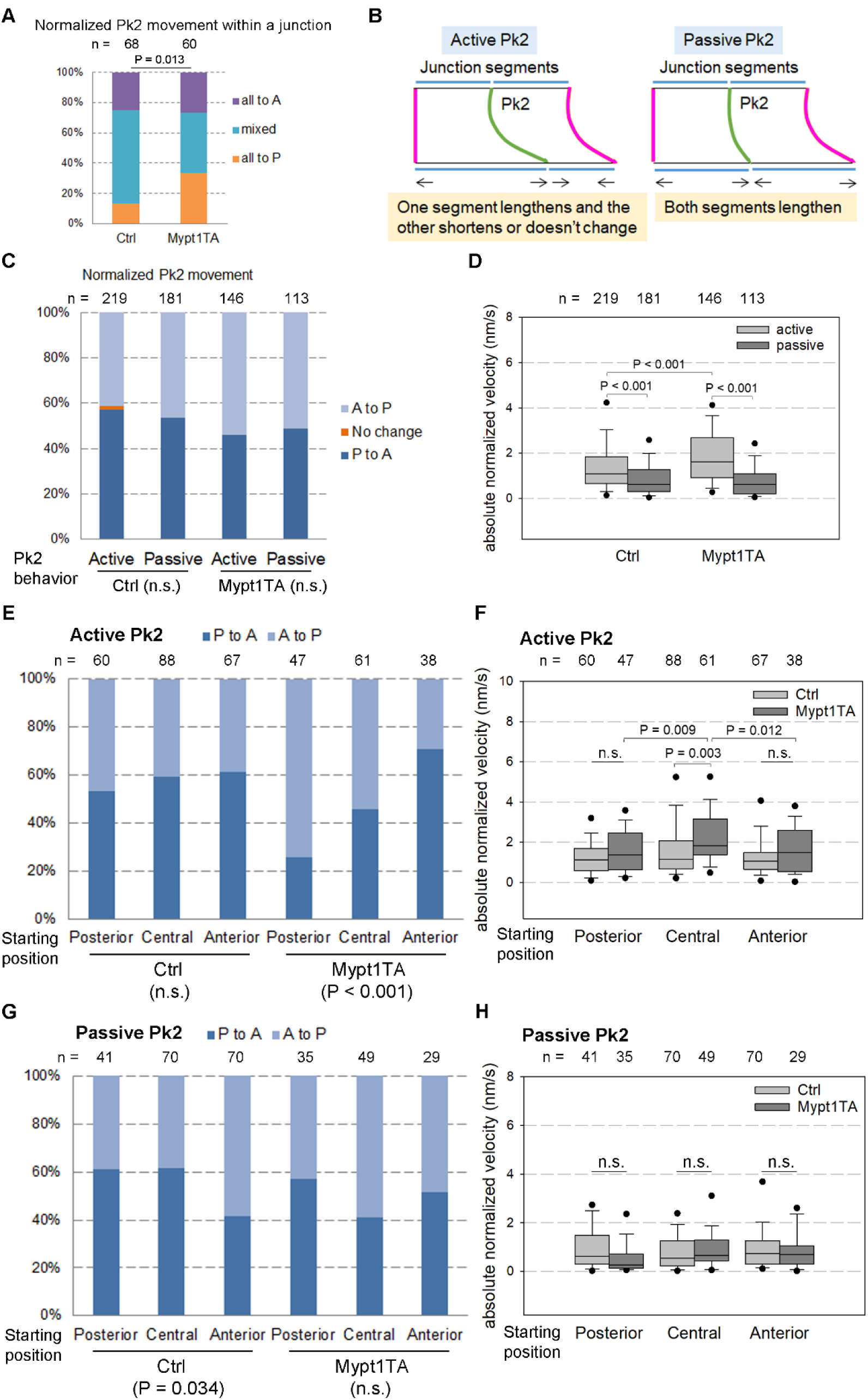
NMII activity regulates the movement of type 1 Pk2 puncta. (A) Percentage of junctions with all their Pk2 puncta moving to the same direction (all to anterior (A) or posterior (P)) or not (mixed). Chi-square test was performed to obtain the *p* values. (B) Schematic illustration of type 1 and type 2 Pk2 puncta behavior at a growing junction. (C) Distribution of normalized Pk2 moving direction based on the behavior of Pk2 puncta in control and Mypt1TA-expressing embryos. (D) Absolute normalized velocity of Pk2 puncta based on their behaviors in control and Mypt1TA-expressing embryos. Two-way ANOVA was performed to obtain the *p* values. (E) Distribution of normalized Pk2 moving direction based on the starting position of type 1 Pk2 puncta at junctions in control and Mypt1TA-expressing embryos. Chi-square test was performed to obtain the *p* values. (F) Absolute normalized velocity of type 1 Pk2 puncta based on their starting position at junctions in control and Mypt1TA-expressing embryos. Two-way ANOVA was performed to obtain the *p* values. (G) Distribution of normalized Pk2 moving direction based on the starting position of type 2 Pk2 puncta at junctions in control and Mypt1TA-expressing embryos. Chi-square test was performed to obtain the *p* values. (H) Absolute normalized velocity of type 2 Pk2 puncta based on their starting position at junctions in control and Mypt1TA-expressing embryos.

Since our data suggests the length of anterior and posterior junction segments flanking Pk2 puncta are independently regulated (Figure 2E), we decided to take a closer look at how Pk2 movement is affected by junction heterogeneity. We began by sorting Pk2 puncta based on how their anterior and posterior junction segments change length (Figure 4B). Pk2 puncta were categorized as “type 1” if the two flanking segments behave differently, for example one is growing while the other is shrinking (Figure 4B, left); conversely, Pk2 puncta were defined as “type 2” if the junction segments flanking them are in the same state, i.e. both are growing or shrinking (Figure 4B, right). We reason that the type 2 category reflects a state of coherent but potentially heterogeneous expansion or shrinkage of the “junctional matrix” that carries the Pk2 puncta, while the type 1 reflects discontinuous junction behavior around the puncta. We next analyzed the frequency of these two modes and found 54% of Pk2 puncta in control cells exhibit the type 1 mode (Figure 4-Figure Supplement 1), indicating that both the type 1 and type 2 modes of transport contribute to Pk2 puncta movement. Interestingly, anterior movement is more common in both the type 1 and type 2 puncta (Figure 4C), implicating processes that bias the direction of Pk2 puncta translation regardless of how the movement is achieved.

We next examined whether NMII activity plays a role in determining type 2 versus type 1 modes of Pk2 puncta motion. Similar to the control cells, the frequency of type 1 Pk2 puncta in Mypt1-expressing cells was 56% (Figure 4- Figure Supplement 1), with a slightly decreased anterior bias in both groups (Figure 4C). As expected, we found the absolute NV of type 1 puncta was higher than that of type 2 ones; but we were surprised to see that Mypt1TA expression specifically increased the absolute NV of the type 1 puncta (Figure 4D). Taken together, these data suggest that NMII does not determine by which mode Pk2 puncta move but plays a more indirect role in the movement of type 1 Pk2 puncta. Following up on this observation, we found that compared to the control cells, Mypt1TA expression led type 1 Pk2 puncta to move toward their nearest vertex (Figure 4E) and increased the absolute NV of type 1 Pk2 puncta within the central region of junctions (Figure 4F). These observations are consistent with our analysis of all the Pk2 puncta (Figure 3), suggesting that NMII regulates directional movement of type 1 Pk2 puncta.

Curious about how the movement of type 2 Pk2 puncta is affected by NMII activity, we also examined the correlation between the starting position of these puncta and their directional motion. We found a weak correlation in control cells that type 2 Pk2 puncta at anterior regions tend to move away from the anterior vertex (Figure 4F, *p* = 0.034) but did not observe this phenomenon in cells expressing Mypt1TA (Figure 4G). Furthermore, there was no correlation between the starting position of type 2 Pk2 puncta and the magnitude of absolute NV (Figure 4H). Taken together, these findings suggest that type 2 Pk2 movement is neither affected by the starting position of the puncta within in the junction, nor by NMII activity.

### NMII regulation of Pk2 puncta movement is more prominent in growing and disturbed junctions

Since Pk2 is reported to be more stable at shrinking junctions than growing junctions (Butler and Wallingford, 2018), we wondered whether junction behaviors and Pk2 puncta movement patterns were correlated. To start, we categorized junction behaviors by their length change over 10 minutes into “idle” (between +1 and −1 μm change), “growing” (greater than 1 μm), and “shrinking” (less than −1 μm); junctions experiencing dramatic but transient length changes were put into the “disturbed” category (Figure 5A). To evaluate the effect of NMII activity, we carefully titrated Mypt1TA mRNA injections so that expression of Mypt1TA didn’t change the distribution of junction behaviors compared to control tissues (Figure 5B). Interestingly, there was no clear correlation between junction behaviors and Pk2 directional movement in control cells; however, Mypt1TA expression significantly reduced the frequency of anteriorly-moving Pk2 puncta within both growing and disturbed junctions (Figure 5C). Moreover, we found that Pk2 puncta in idle junctions had significantly lower absolute NV compared to those within other types of junctions, and expression of Mypt1TA didn’t change this correlation even though the absolute NV of puncta within idle junctions increased (Figure 5D). Together these results suggest that the speed of Pk2 puncta motion correlates with junction length changes and that NMII activity is specifically required to bias the direction of Pk2 movement in growing and disturbed junctions, categories that both show length increases.

**Figure 5.**
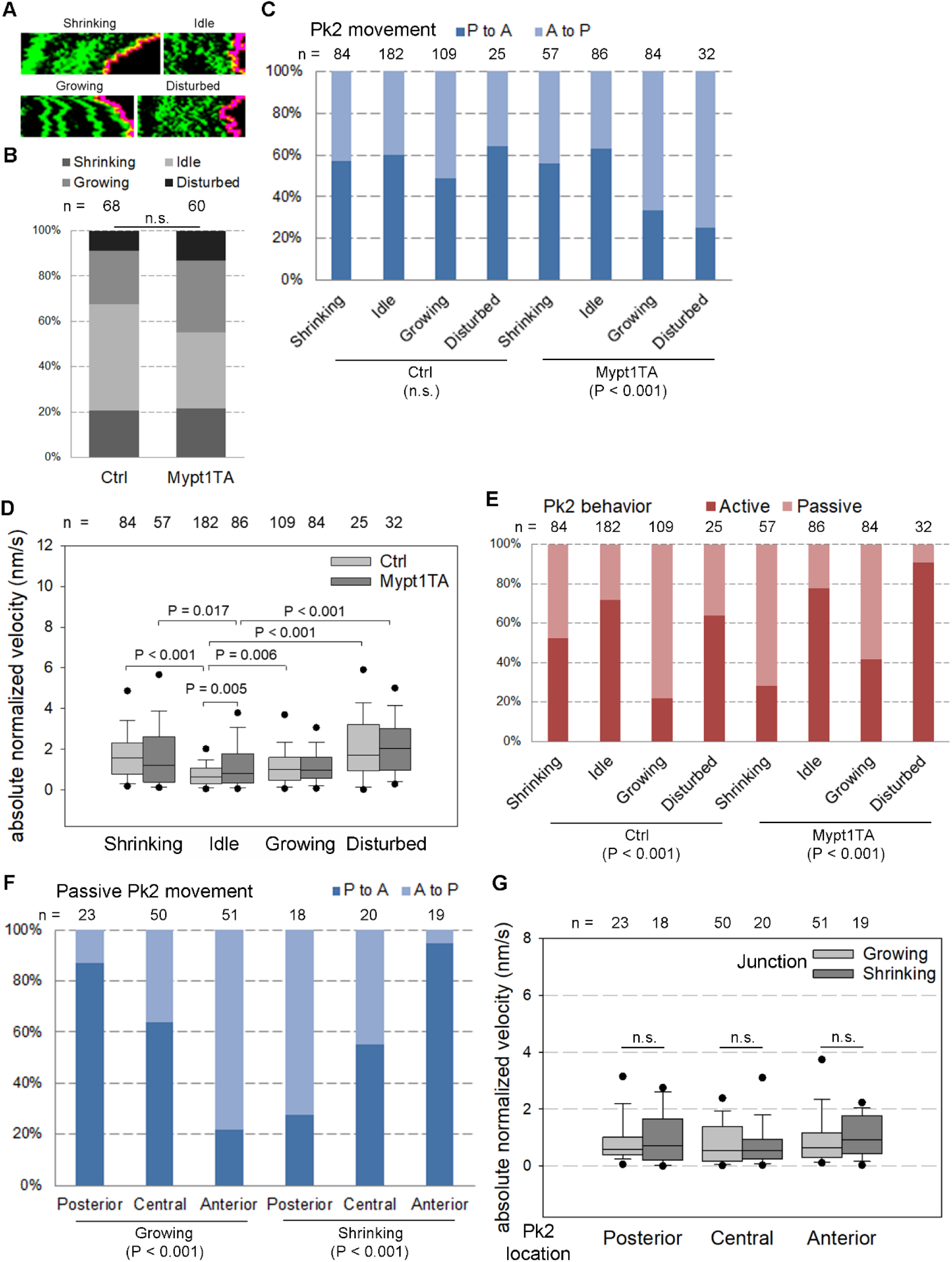
Correlations between the junction behavior and Pk2 movement. (A) Representative kymographs of the indicated junction behavior distinguished by the trace of the right junction end (red). (B) Distribution of junction behaviors in control and Mypt1TA-expressing embryos. n, number of junctions. Chi-square test was performed to obtain the *p* values. (C) Distribution of normalized Pk2 moving direction based on junction behaviors in control and Mypt1TA-expressing embryos. n, number of Pk2 puncta. Chi-square test was performed to obtain the *p* values. (D) Absolute normalized velocity of Pk2 puncta based on junction behaviors in control and Mypt1TA-expressing embryos. Two-way ANOVA was performed to obtain the *p* values. (E) Distribution of Pk2 behaviors based on junction behaviors in control and Mypt1TA-expressing embryos. Chi-square test was performed to obtain the *p* values. (F) Distribution of normalized Pk2 moving direction based on the starting position of type 2 Pk2 puncta at growing and shrinking junctions. Chi-square test was performed to obtain the *p* values. (G) Absolute normalized velocity of type 2 Pk2 puncta based on their starting position at growing and shrinking junctions. Two-way ANOVA was performed to obtain the *p* values.

Given that NMII activity regulates the movement of type 1 Pk2 puncta, we next examined the distribution of type 1 or type 2 Pk2 puncta in idle, growing, shrinking or disturbed junctions. Considering that idle junctions by definition had almost no change of whole junction length over time, we expected that most Pk2 puncta within idle junctions would be grouped as type 1 and it was indeed the case (Figure 5E). Interestingly, growing junctions have the highest percentage of type 2 Pk2 puncta, suggesting that junction growth events are distributed evenly along the junction compared to junction shrinkage events. Intriguingly, expression of Mypt1TA increased the ratio of type 1 Pk2 puncta specifically in growing and disturbed junctions (Figure 5E), suggesting that NMII activity plays a role in the even distribution of junction growth events along the length of each junction. In summary, we find that Pk2 puncta preferentially move via the type 2 process in growing junctions and that NMII activity regulates this process.

### Analysis of type 2 Pk2 movement reveals that junction length change is most prominent at junction ends

Since most Pk2 puncta in growing junctions are type 2 but more mobile than those in idle junctions, we concluded that junction growth usually happens throughout the junction but with differential local growth rates, driving type 2 Pk2 movement. To better understand how junction growth is spatially regulated within individual junctions, we analyzed the correlation between the starting position of type 2 Pk2 puncta in growing junctions and their directional motion. Intriguingly, type 2 puncta in growing junctions tend to move away from the nearest vertex (Figure 5F, growing, *p* < 0.001), indicating that junction growth is faster near vertices. Curious about what would be the case at shrinking junctions, we carried out the same analysis for type 2 Pk2 puncta within shrinking junctions. Contrary to the behavior of type 2 puncta in growing junctions, those in shrinking junctions move toward the nearest vertex (Figure 5F, shrinking, *p* < 0.001), consistent with the idea that junctions shrink faster at or near the vertices. Moreover, the direction of Pk2 puncta movement, but not their speed, correlates to their starting position within growing or shrinking junctions (Figure 5G). Furthermore, Mypt1TA expression did not change the direction of Pk2 puncta movement (Figure 5- Figure Supplement 1), indicating that NMII activity plays little or no role in the directional bias of type 2 puncta motion. Taken together, our analyses of type 2 Pk2 movement suggest that junction growth or shrinkage occurs near vertices in an NMII-independent manner.

### Pk2 partitions together with ZO-1

If Pk2 puncta movement is due to heterogeneous junction growth and shrinkage, it is expected that 1) the apical junctions of presumptive neuroectoderm exhibit mechanical heterogeneity, and 2) other junctional proteins proximal to Pk2 puncta move the same way as Pk2. To test whether the junctions are compartmentalized by cadherin clusters, we assessed the distribution of E-cadherin, the major cadherin localized to the apical junction by the end of gastrulation and throughout neurulation (Nandadasa et al., 2009; Session et al., 2016; Van Itallie et al., 2025). Compared to the membrane marker, E-cadherin tagged with mNG formed visible puncta at the apical junctions (Figure 6A-A’). The “punctaness” of the junctional signals were then calculated based on the shape metrics representing smoothness, variability, and puncta, and we found that E-cadherin exhibited higher “punctaness index” than the membrane marker (Figure 6B). These results suggest that the apical junctions contain clusters of E-cadherin and are possibly mechanically heterogeneous.

**Figure 6.**
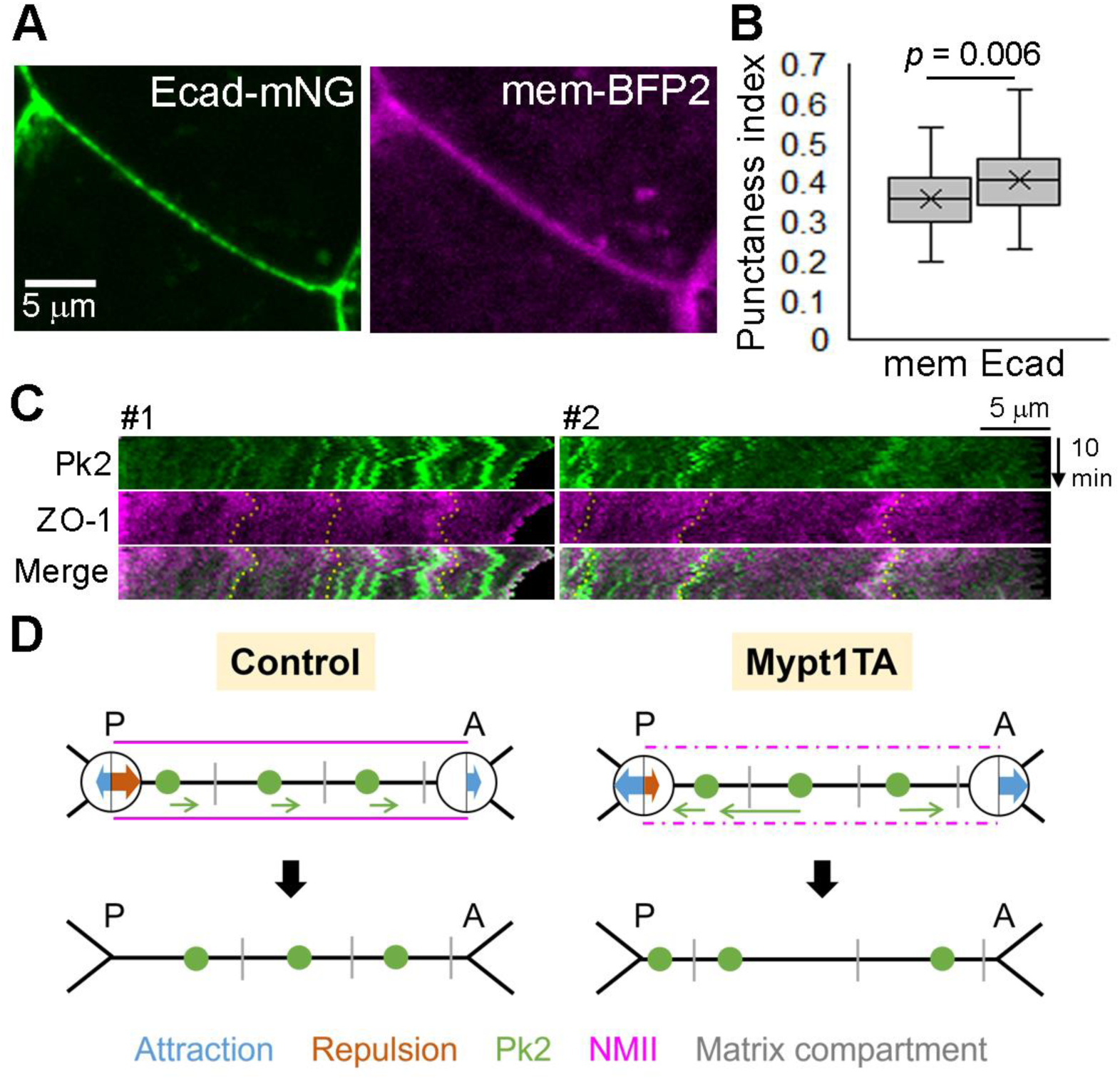
ZO-1 partitioned together with Pk2. (A) Representative images of an apical junction of stage 12 presumptive neuroectoderm expressing E-cadherin-mNG (green) and mem-mTagBFP2 (magenta). (B) Box plot showing the punctaness index of the junctional signal of mem-mTagBFP2 (mem) and E-cadherin (Ecad). Data are from 57 junctions of 2 embryos. Paired Student’s t-test was performed to obtain the *p* values. (C) Two representative kymographs showing the displacement of mNG-Pk2 and RFP-ZO-1 along the bicellular junction. Dotted lines depict the right-side edge of the ZO-1 patches. (D) Working model. Pk2 puncta (green) movement along a junction is driven by repulsion (brown arrow) from the posterior (P) vertex due to biased expansion of the posterior junction segment, leading to Pk2 movement toward the anterior (A) vertex. Peri-junctional NMII (magenta) compartmentalization (grey) within the apical junctional matrix and restricts the magnitude of Pk2 displacement. Reducing NMII activity by expressing Mypt1TA weakens compartmentalization, leading to overall less repulsion and more unrestricted Pk2 movement.

We next sought to test if other junctional proteins show similar moving patterns to that of Pk2 puncta. To avoid potential interference between fluorescently-tagged Pk2 and adherens junction proteins, we evaluated the movement of ZO-1, a tight junction protein (Stevenson et al., 1986), along the bicellular junction in the presence of Pk2. The naturally uneven distribution of ZO-1 presented some traceable edges of the ZO-1 patches, which moved similarly to their nearby Pk2 puncta (Figure 6C). These findings suggest that the Pk2 movement we observed reflects junction heterogeneity that affects the displacement of junctional proteins in general.

## Discussion

In this study, we investigated junctional Pk2 puncta movement in *Xenopus* gastrula neuroectoderm by an in-depth analysis of junction growth and dynamics. We uncovered that Pk2 puncta movements are biased toward the anterior-most vertex and are regulated by NMII. The observed low speed of Pk2 movement suggests that movements of Pk2 puncta reflect complex, spatially heterogeneous processes that contribute to changes of junction length. Moreover, NMII activity is required to reduce nonselective Pk2 movements toward cell vertices and enhance anterior movement bias. We propose that intra-junctional heterogeneity, i.e. variation in the composition and biophysical properties of the apical junctional matrix, contributes to PCP by partitioning Pk2 toward the anterior vertex and that this process is one mechanism by which NMII activity regulates PCP.

Our findings revealed that the anterior-ward Pk2 puncta movements within apical junction segments are driven by the “repulsion” of Pk2 from the posterior vertex due to biased expansion of the posterior junction segment (Figure 6B, brown arrow). Analysis of type 2 Pk2 puncta movement suggests that both junction growth and shrinkage occur via junctional matrix being added or removed near each vertex, and we propose that these mechanisms are responsible for the observations that each vertex can either “repel” or “attract” puncta. It is important to note that the anterior-ward bias of puncta movement is small, and that accumulation of significant levels of Pk2 at anterior cell interfaces is likely due to repeated cycles of junctional matrix addition.

Our observation of the biased length change of the junction segments flanking Pk2 puncta parallels recent reports that bicellular junctions are physically heterogeneous irrespective of their role in PCP signaling (Huebner et al., 2021; Mei et al., 2014). The precise mechanisms regulating junctional heterogeneity appear to include spatial and temporal regulation of cell-cell adhesions, membrane trafficking, and cytoskeleton. Cis-clustering of cadherins as well as intercellular trans-dimers (Yap et al., 2015) have been proposed to regulate cytoskeletal compartmentalization at the lateral cortex and adherens junctions (Krapf, 2018). Indeed, two recent studies suggest junctional heterogeneity reflects fluctuations of C-cadherin activity along mesenchymal cell-cell contacts (Huebner et al., 2021) as well as NMII-mediated contractility along the adherens junctions (Yang et al., 2022). Moreover, NMII position within the cell cortex, either near to or more distant from the plasma membrane, regulates mechanical properties of the cortex and allow rapid solid-to-fluid transitions in cell surface mechanics (Truong Quang et al., 2021). Since PCP complexes are localized in neuroectoderm at the level of adherens junction (Butler and Wallingford, 2017; Shimada et al., 2001), we propose that intra-junctional mechanical heterogeneity is coordinated through NMII activity and cadherin clustering.

What molecular mechanisms are responsible for junction heterogeneity and directed motion of Pk2? As these processes are likely confined to regions near the posterior vertex, it is possible that PCP complexes are compartmentalized at posterior junctions by cadherin clusters, septins, or other components of the junctional matrix (Huebner et al., 2021; Shindo and Wallingford, 2014) that initiate a positive feedback of directional displacement. The displacement may be due to heterogeneous addition or removal of the plasma membrane or contraction of the perijunctional actomyosin belt. It is important to recognize that the dynamics of apical junctional complexes are likely dissimilar to the dynamics of the low-adhesion sites and associated cell cortex at basolateral along cell-cell interfaces in sheets of mesenchymal cells, such as those involved in mesoderm convergent extension (see (Green and Davidson, 2007)). Consistent with this idea, we observed E-cadherin clusters at the junction of presumptive neuroectoderm. It remains an open question as to whether and how cadherins contribute to Pk2 movement along the junctions.

While we were unable to analyze E-cadherin and Pk2 displacement together due to technical reasons, we found that ZO-1 patches partition together with Pk2, supporting the hypothesis that the protein displacement is due to junction heterogeneity and not specific to Pk2. Notably, ZO-1 is located at the tight junction but still partitions alongside Pk2, suggesting that they are carried by the junction matrix that couples the tight junction and adherens junction components. Consistent with this idea, ZO-1 is known to scaffold junctional complexes by interacting with adherens junction components such as α-catenin and Afadin (Ooshio et al., 2010; Rajasekaran et al., 1996). Importantly, ZO-1 partitions with Pk2 but does not become anteriorly polarized, possibly due to its high turnover rate (Beutel et al., 2019; Schwayer et al., 2019; Shen et al., 2008) as compared to the increasingly stabilized Vangl-Pk complexes during polarization (Nissen et al., 2026; Strutt et al., 2011). While ZO-1 has been used as fiducial position markers alongside local RhoA activation (Stephenson et al., 2019), it remains to be seen whether ZO-1 is involved in establishing the boundary of the junction compartments.

NMII activity is a prerequisite for PCP establishment (Ossipova et al., 2015; Winter et al., 2001) as well as a downstream target of PCP signaling pathway (Habas et al., 2003; Kim and Davidson, 2011; McGreevy et al., 2015; Tahinci and Symes, 2003). We assessed roles of NMII activity in junctional Pk2 movement and found that NMII regulates Pk2 movement in two aspects (Figure 6B): 1) maintaining a “repulsion” from the posterior segment, and 2) reducing Pk2 moving speed. It is known that NMII plays critical roles in adherens junction assembly, including bundling peri-junctional F-actin filaments and connecting cadherin complexes to F-actin bundles (Choi et al., 2016; Heuzé et al., 2019; Shewan et al., 2005; Smutny et al., 2010; Yu-Kemp et al., 2022). In addition, actomyosin cytoskeletons have been shown to organize membrane compartments and restrict protein diffusion in vitro and in vivo (Goswami et al., 2008; Köster et al., 2016; Vogel et al., 2017). We propose that NMII restricts junctional Pk2 movement by maintaining apical cortex compartmentalization. Actomyosin contractility and adhesion together could operate as a “mechanical clutch” that physically couples or traps Pk2 puncta within the junctional matrix. Our finding that most Pk2 puncta move by sub-diffusion also supports this hypothesis. Notably, we find shrinking junctions exhibit more intra-junctional heterogeneity (i.e. a higher percentage of type 1 Pk2 puncta) compared to growing junctions, perhaps indicating NMII engagement in the “clutch”. Moreover, NMII might contribute to the asymmetric repulsion of Pk2 puncta from posterior vertices by promoting mechanical heterogeneity within the junction, perhaps through the transient accumulation of cadherins (Cavanaugh et al., 2022; Vanderleest et al., 2018) or by spatial control of fluid-solid transitions in the cortical actomyosin cytoskeleton (Truong Quang et al., 2021). Taken together, we propose that NMII regulates Pk2 puncta movement by fine-tuning the distribution of adhesion complexes or by regulating cytoskeletal viscoelasticity along the junction. It remains to be tested whether NMII plays a general role in partitioning junctional proteins.

One interesting observation is that reducing NMII activity leads to Pk2 moving toward both the anterior and posterior vertices, implying that this moving pattern may reflect the “default” junction heterogeneity. We did not observe Pk2 accumulation at the vertices as a result of this phenotype, which could be due to higher Pk2 turnover rate at the vertices and/or bicellular junctions. The idea that NMII activity regulates Pk2 turnover is consistent with the correlation between Pk2 and Myl9 dynamics during neuroepithelial junction shrinking (Butler and Wallingford, 2018).

In summary, our data support a hypothesis that Pk2 polarization reflects intra-junctional heterogeneity and association with the junctional matrix that is maintained by NMII activity. Furthermore, it is clear from our work that Pk2 puncta do not move toward anterior sites as a conventional motor-bound cargo. We identified the “type 1” and “type 2” mode of Pk2 movement associated with higher or lower junction heterogeneity, respectively. Interestingly, reducing NMII activity had more impact on the type 1 mode but did not convert them into type 2, suggesting that the type 1 mode involves other unidentified factors in addition to the higher NMII activity. Future work that combines live imaging of NMII, apical junctional complexes, and Pk2 is needed to evaluate mechanisms that integrate adhesion and NMII into a mechanical-clutch, whether such a system plays a unique role in segregating Pk2 to the anterior junctions in neuroectoderm, and how these systems mediate directed cell rearrangement during convergent extension.

## Acknowledgement

We thank Sommer Anjum for advice on the image analysis macro, John Wallingford for the Pk2 construct, Sang-Wook Cha for the E-cadherin construct, and Ann Miller for the SF9 and ZO-1 constructs. We are grateful to the Davidson lab members for discussions. CC and LAD were supported by a grant from the NIH to LAD (R01 HD044750).

## Materials and Methods

### DNA constructs and *Xenopus laevis* embryo manipulation

The mNeonGreen-Pk2 and E-cadherin-mNeonGreen constructs were made by subcloning mNeonGreen and *Xenopus* Pk2 or E-cadherin (Cdh1) into pCS2+ (Butler and Wallingford, 2015; Nandadasa et al., 2009; Shaner et al., 2013), and the Mypt1TA (aka MBS^T695A^; (Jackson et al., 2017)) construct is subcloned into pCS2+ from Addgene plasmid 31669 (Banko et al., 2011). The RFP-ZO-1 and SF9-mNeonGreen constructs are gifts from Ann Miller (Arnold et al., 2019; Stephenson et al., 2019).

All *Xenopus* laevis frogs used in this study were wild type pigmented and obtained commercially (Nasco, Fort Atkinson, WI). Embryos were obtained by in vitro fertilization following standard protocols (Sive et al., 2000). Fertilized embryos were de-jellied in 2% cysteine solution (pH 8) and staged accordingly (Nieuwkoop and Faber, 1994). 100 pg of RNAs were microinjected dorsal animally into embryos at four-cell stage, and embryos were cultured in 1/3 X modified Barth’s Solution (MBS) until mid-gastrula stage (stage 11).

### Live imaging and image quantification methods

After manually removing vitelline membranes from the embryos, 180° marginal zone explants were excised and mounted in a custom imaging chamber filled with Danilchik’s For Amy (DFA) medium (Chu and Davidson, 2021; Green et al., 2004). Explants were allowed to heal for 10 minutes, and time-lapse images were acquired using a custom-built spinning disk confocal microscope with a 63X oil immersion objective. Images were taken every 30 seconds for 10 minutes using Micro-Manager (Edelstein et al., 2014), and max intensity projection images were created from 3-5 μm z stacks of images.

Images of 4 explants from 2 clutches (2 explants per clutch) were processed using Fiji (ImageJ 1.53, NIH), and Tissue Analyzer was used to segment the cells and produce kymographs of each boundary (Aigouy et al., 2010). For tracking trajectories, we wrote a custom macro to perform z-score normalization of the Pk2 signals to reduce the background and highlight the right edge of the kymograph based on the mCherry signals. Trajectories of Pk2 puncta and the junction right end were tracked using KymoButler for ImageJ (Jakobs et al., 2019) and manual correction in case that visible mis-connection happened. Another custom macro was used to calculate the velocity and normalized velocity of each trajectory, and the velocity of the left junction end naturally becomes 0 due to kymographs being aligned to the left end. Junction orientation data collected by Tissue Analyzer was used to determine the anterior and posterior junction ends. Velocity was defined as particle displacement over time, and normalized velocity was defined as changes of the particle relative position within the junction over time. An anterior-moving particle is defined as a particle with positive value of normalized velocity, and absolute velocity or normalized velocity is defined as the absolute value of the velocity or normalized velocity respectively. The anterior or posterior segments were defined as the junction segments anterior or posterior to the particle respectively, and segment length change was defined as the velocity difference between the particle and the corresponding junction end. To better assess Pk2 movement pattern within each junction, only junctions with 2 or more Pk2 puncta 1 μm away from junction ends were used. To quantify the clustering of E-cadherin, we computed the coefficient of variation, RMS of the intensity roughness, and peak counts and prominences of E-cadherin signals at apical junctions. These parameters are linearly combined into a “punctateness index” with high values indicating punctate patterns and low values indicating smooth patterns. Data were plotted using Excel (Microsoft, Redmond, WA) and SigmaPlot (Systat Software, Chicago, IL), and statistical analyses were performed with SigmaPlot. Unless otherwise noted, all box plots show the 25th, median, and 75th percentiles of the data. The error bars represent the 10^th^ and 90^th^ percentiles, and the dots mark the 5^th^ and 95^th^ percentiles.

---

**Figure 1- Figure Supplement 1.**
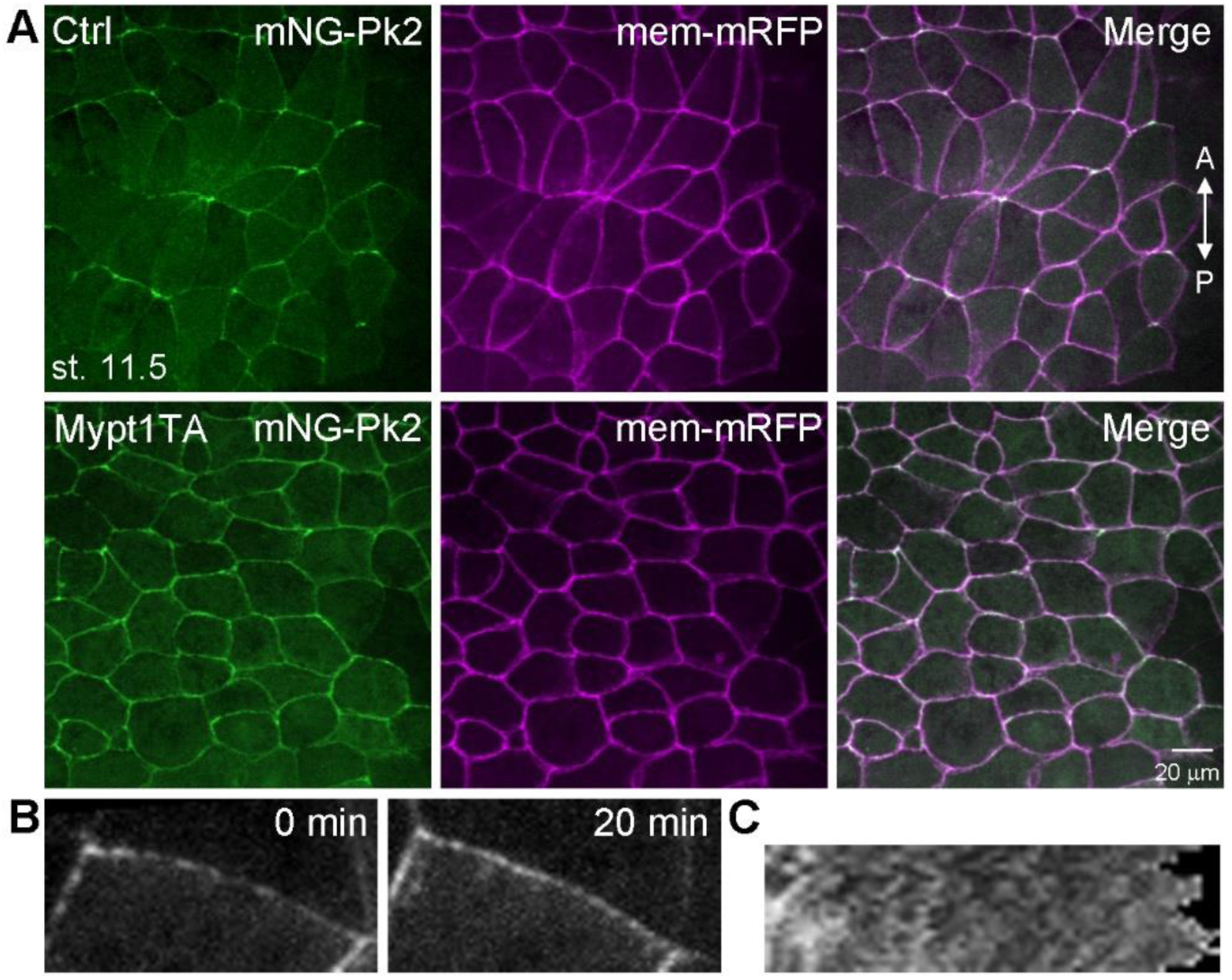
Mypt1TA expressing abolished Pk2 polarization. (A) Embryos injected with mRNAs encoding indicated proteins were fixed at stage 11.5, and epifluorescence of presumptive neuroectoderm was shown. Note the enrichment of mNG-Pk2 at junctions orthogonal to the AP axis in control (Ctrl) but not Mypt1TA-expressing embryos. (B) Stills of a representative movie of junctional mNG-Pk2 in the presence of Mypt1TA at the indicated time points. (C) Kymograph of the junction in (B).

**Figure 2- Figure Supplement 1.**
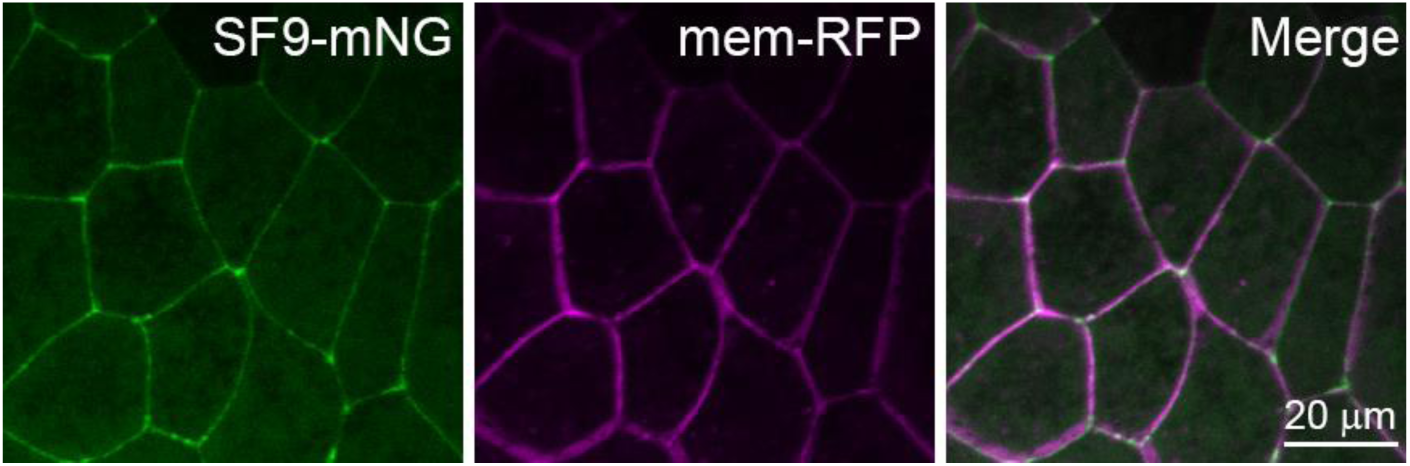
Distribution of SF9 at the apical junctions. Embryos injected with mRNAs encoding indicated proteins were fixed at stage 11, and epifluorescence of presumptive neuroectoderm was shown.

**Figure 2- Figure Supplement 2.**
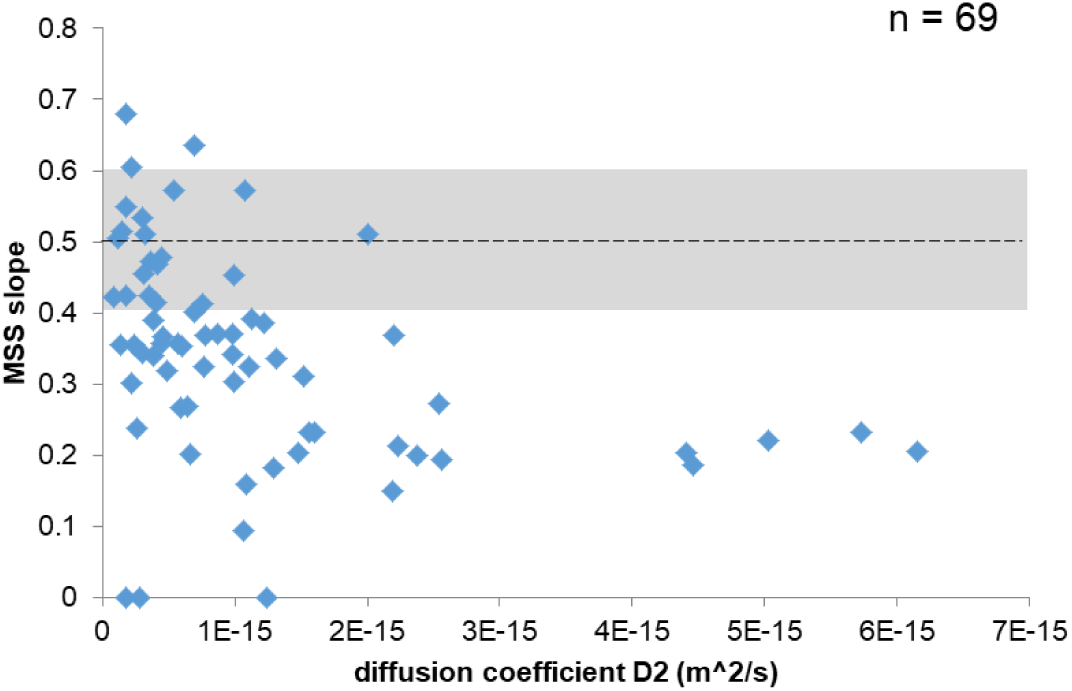
Scatter plot showing the distribution of diffusion coefficient and MSS slope of junctional Pk2 puncta from a time-lapse movie. The MSS slope of a pure diffusion event equals to 0.5, and slopes above and below 0.5 represent super-diffusion and sub-diffusion, respectively.

**Figure 2- Figure Supplement 3.**
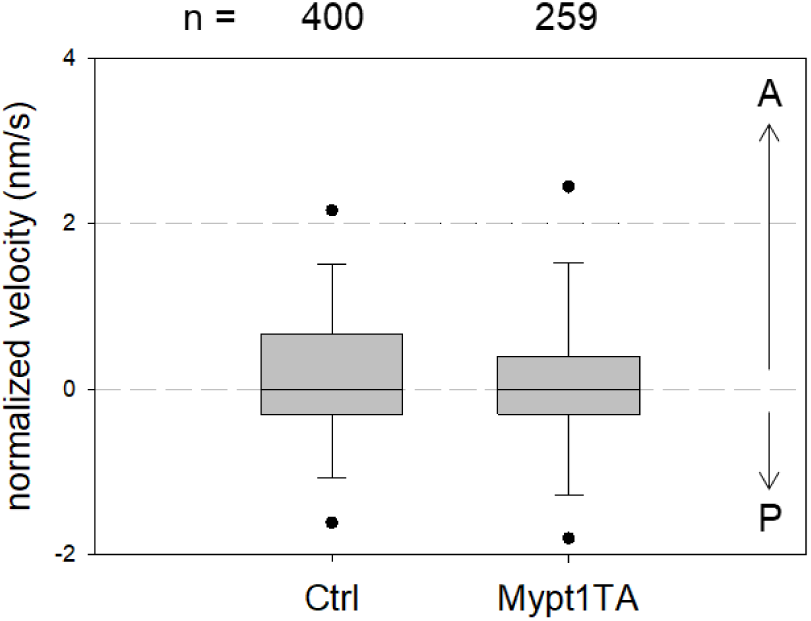
Normalized velocity of Pk2 puncta in control and Mypt1TA-expressing embryos. A positive value of velocity means that Pk2 moves toward the anterior end of junction. n, number of Pk2 puncta.

**Figure 4- Figure Supplement 1.**
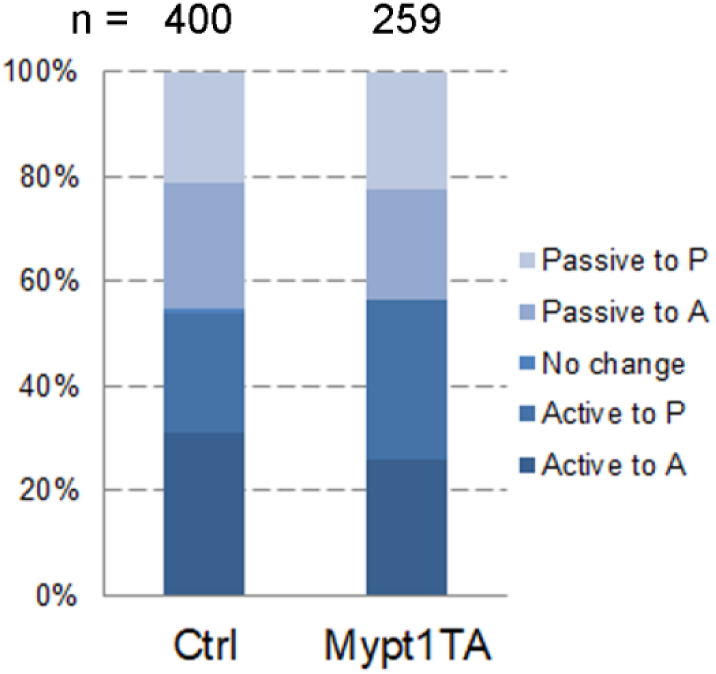
Distribution of Pk2 behaviors and normalized moving direction in control and Mypt1TA-expressing embryos.

**Figure 5- Figure Supplement 1.**
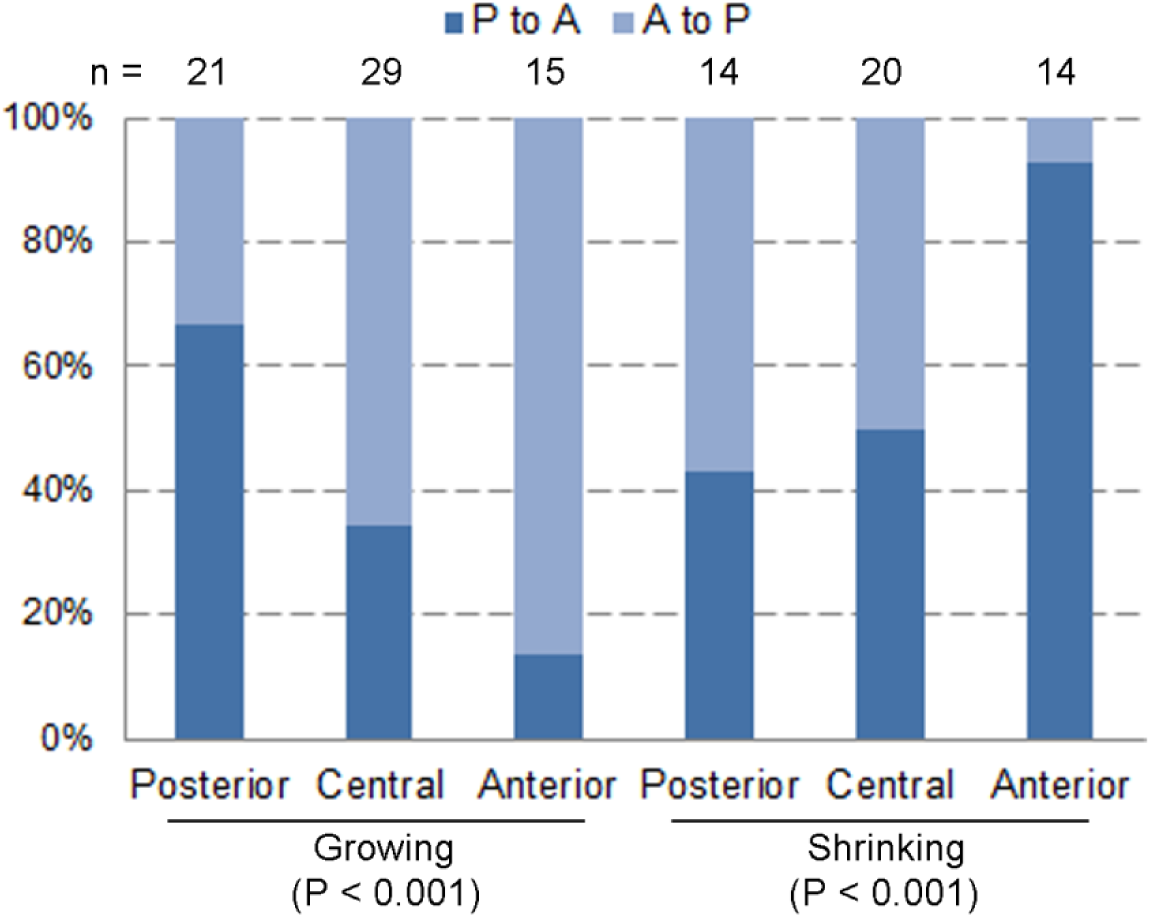
Distribution of normalized Pk2 moving direction based on the starting position of type 2 Pk2 puncta at growing and shrinking junctions in Mypt1TA-expressing embryos. Chi-square test was performed to obtain the *p* values.

## Supplemental Materials and Methods

### MSS analysis of Pk2 movement

The x-y coordinates of Pk2 traces were imported into Mosaic Particle Tracker (MosaicSuite for Fiji) (Sbalzarini and Koumoutsakos, 2005), and the mean square displacement (MSD) and slope of moment scaling spectrum (MSS) were calculated for each trace.

### Supplemental Movies

Supplemental movie 1 (related to Figure 1). Time lapse images of the presumptive neuroectoderm of embryos expressing mNG-Pk2 at stage 11. Note the movement of junctional Pk2 puncta toward the vertices.

Supplemental movie 2 (related to Figure 1). Time lapse images of the presumptive neuroectoderm of embryos expressing mNG-Pk2 and Mypt1TA at stage 11. Note the lack of Pk2 polarization despite Pk2 puncta movement toward the vertices.

